# Plasticity in *Escherichia coli* cell wall metabolism promotes fitness and mediates intrinsic antibiotic resistance across environmental conditions

**DOI:** 10.1101/379271

**Authors:** Elizabeth A Mueller, Petra Anne Levin

## Abstract

Although the peptidoglycan cell wall is an essential structural and morphological feature of most bacterial cells, the extracytoplasmic enzymes involved in its synthesis are frequently dispensable under standard culture conditions. By modulating a single growth parameter—extracellular pH—we discovered a subset of these so-called “redundant” enzymes in *Escherichia coli* are required for maximal fitness across pH environments. Among these pH specialists are the class A penicillin binding proteins PBP1 a and PBP1 b; defects in these enzymes attenuate growth in alkaline and acidic conditions, respectively. Genetic, biochemical, and cytological studies demonstrate that synthase activity is required for cell wall integrity across a wide pH range, and differential activity across pH environments significantly alters intrinsic resistance to cell wall active antibiotics. Together, our findings reveal previously thought to be redundant enzymes are instead specialized for distinct environmental niches, thereby ensuring robust growth and cell wall integrity in a wide range of conditions.

## INTRODUCTION

The growth and survival of single-celled organisms relies on their ability to adapt to rapidly changing environmental conditions. A commensal, pathogen, and environmental contaminant, *Escherichia coli* occupies and grows in diverse environmental niches, including the gastrointestinal tract, urinary bladder, freshwater, and soil. In the laboratory, the bacterium’s flexibility in growth requirements is reflected in robust proliferation across a wide range of temperature, salt, osmotic, pH, oxygenation, and nutrient conditions (1).

The physiological adaptations that permit growth and survival across environmental conditions are not yet well understood, particularly for extracytoplasmic processes. Due to the discrepancy in permeability between the plasma and outer membrane (2), the periplasmic space of Gram negative bacteria is highly sensitive to chemical and physical perturbations, including changes in salt, ionic strength, osmolality, and pH. Notably, upon mild environmental acidification, the periplasm assumes the pH of the extracellular media, while the cytoplasmic pH remains minimally disrupted (3,4). Although mechanisms that contribute to cytoplasmic pH homeostasis have been described (5,6), little is known about the quality control mechanisms that preserve proper folding, stability, and activity of key proteins in the periplasm in the absence of a homeostatic control system.

The peptidoglycan (PG) cell wall and its synthetic machinery are among the key constituents of the periplasm that must be preserved across growth conditions. Essential for viability among most bacteria, PG is composed of glycan strands of repeating *N-* acetylglucosamine and N-acetylmuramic acid disaccharide units crosslinked at peptide stems (7). Beyond providing the force necessary to resist turgor pressure, the cell wall reinforces cell shape and serves as a major interface for cell-cell and cell-host interactions (8,9). As an essential process, cell wall synthesis is also the principle target of several classes of antibacterial agents, including β-lactam (e.g. penicillin) and glycopeptide (e.g. vancomycin) antibiotics.

PG precursors are assembled in the cytosol and translocated across the inner membrane into the periplasm, where cell wall synthases construct the PG network through a series of glycosyltransferase (glycan polymerizing) and transpeptidase (peptide crosslinking) reactions. PG synthases include bifunctional class A penicillin binding proteins (aPBPs), as well monofunctional transpeptidases (class B PBPs; bPBPs) and monofunctional glycosyltransferases, typically of the shape, elongation, division, and sporulation (SEDS) protein family (10-12). L,D-transpeptidases synthesize non-canonical 3-3 crosslinks between peptide stems and are predominately active during PG remodeling during stationary phase growth in *E. coli* (13,14). In addition to synthases, a series of periplasmic cell wall hydrolases and autolysins—including D,D-carboxypeptidases, D,D and L,D-endopeptidases, lytic transglycosylases, and amidases—are required to accommodate nascent strand insertion for expansion of the PG network, create substrate binding sites, and separate cells during the final stages of cytokinesis (8). These enzymes may also play a role activating synthases to ensure cell wall integrity (15).

Curiously, high levels of redundancy in periplasmic cell wall enzymes have been observed across bacterial species. In *E. coli* over 36 enzymes carry out the nine reactions that occur in the periplasm, yet the cytoplasmic steps of PG precursor biogenesis have nearly a 1:1 stochiometric ratio (12 reactions: 14 enzymes) (16). Moreover, with the exception of SEDS glycosyltransferases/bPBP pairs of RodA/PBP2 and FtsW/PBP3 required for lateral expansion of the cell wall and cell division, respectively (11,12), the remaining periplasmic cell wall enzymes are nonessential. Inactivation of an individual enzyme—and in some cases, even multiple enzymes in the same class—often fails to confer discernable growth or morphological phenotypes under standard culture conditions (17-21). Although technological breakthroughs have aided in the identification of cell wall metabolic genes encoding proteins with similar functions (e.g. (22)), elucidating the potential fitness benefit to redundancy has proven challenging.

One model to account for the apparent redundancy of periplasmic cell wall proteins is that enzymes within a given class may be specialists for distinct environmental niches, thereby allowing bacteria to cope with the diverse chemical and physical properties that might affect protein stability and function in this compartment (16). In support of this hypothesis, several groups have identified cell wall enzymes that have increased activity in acidic media. Peters and colleagues demonstrated that *E. coli* carboxypeptidase PBP6b plays a key role in maintenance of cell morphology during growth at pH 5.0 (23), while Castanheira *et al*. identified a PBP3 homolog in *Salmonella* Typhimurium that is preferentially involved in septation at low pH (24). Similarly, the lytic transglycosylase MltA exhibits maximal activity in acidic conditions *in vitro* (25), although whether this property is relevant *in vivo* remains unknown.

In light of these findings, we hypothesized that loss of an enzyme specialized for a particular environmental niche may produce a condition-specific growth defect through impaired cell wall integrity, allowing us to take a systems-level approach to identifying enzymes with differential roles in growth *in vivo*. In screening 32 mutants across six classes of nonessential periplasmic cell wall enzymes, we determined that a subset of these enzymes is differentially required for fitness across pH environments. We find that disruptions in in the activity of cell wall synthases PBP1a and PBP1b conferred fitness defects in opposing pH ranges that can be attributed in part to pH-dependent differences in activity. Concerningly, synthase specialization has consequences for intrinsic resistance to β-lactam antibiotics in nonstandard growth conditions.

## RESULTS

### Identification of pH specialist cell wall synthases and hydrolases

To determine the contribution of individual cell wall enzymes to pH-dependent growth, we cultured strains harboring insertional deletions in each of three aPBPs, six L,D-transpeptidases, five carboxypeptidases, four amidases, nine lytic glycosyltransferases, and six endopeptidases to mid-exponential phase (OD_600_ ~0.2-0.6) in buffered LB media (pH 6.9) then sub-cultured them into fresh LB buffered to pH 4.8, 6.9, or 8.2 for growth rate analysis. These pH values were chosen as representative, physiologically relevant conditions *E. coli* cells encounter in the lower GI tract (pH 5-9) or urine (pH 4.5-8) (26,27). Classification as a pH-sensitive mutant required a significant, > 5% decrease in early exponential phase mass doubling time (unpaired t-test, p < 0.01) in at least one pH condition compared to the parental strain.

Collectively, seven mutants displayed significant reductions in mass doubling times (MDT) at one or more pH values. We observed both acid-sensitive and alkaline sensitive mutants across three enzymatic classes. Strikingly, loss of the bifunctional synthase PBP1b *(mrcB)* abolished growth at pH 4.8 but maintained wild type MDT at neutral and alkaline pH (Fig. 1A). Four additional mutants exhibited a distinct, albeit less severe, defect in MDT (6-14% reduction) at pH 4.8, including strains defective for lytic transglycosylases MltA and MltB and endopeptidase MepS *(spr)* (Fig. 1B-C). Consistent with roles in acid tolerance, MltA was previously shown to have elevated enzymatic activity in acidic conditions *in vitro* (25), and MltB was recently found to contribute to acid shock survival in *Acinetobacter baumannii* (28).

**Figure 1.**
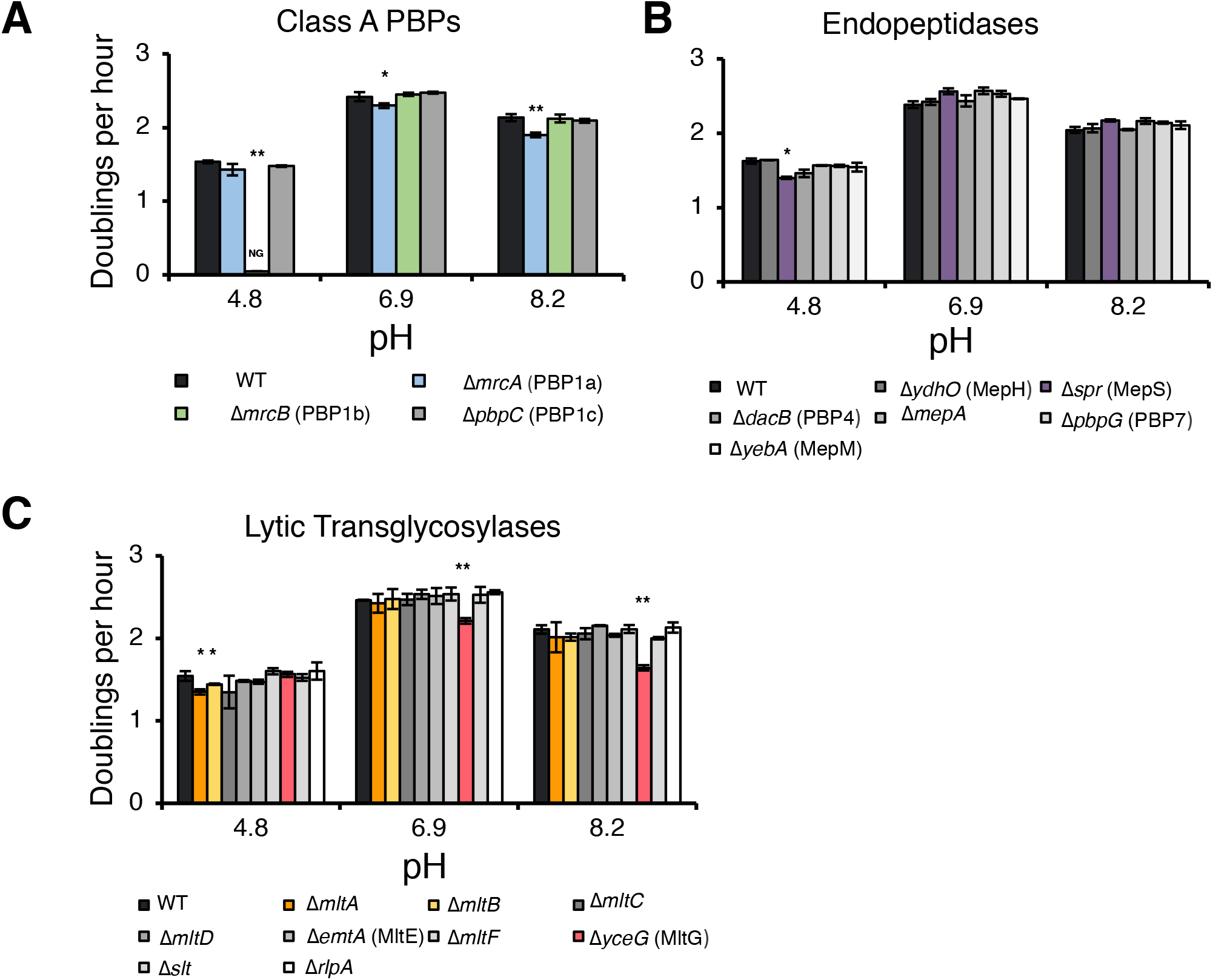
Identification of pH specialist cell wall enzymes. Insertional deletions in non-essential class A PBPs (A), endopeptidases (B), and lytic transglycosylases (C) were screened for growth defects compared to the parental strain in LB media buffered to pH 4.8, 6.9, and 8.2. Bars depict average doublings per hour +/- standard deviation of three independent biological replicates. NG denotes “no growth” observed throughout the course of the experiment (20 hours). Asterisks denote significance of strains with a < 5% growth defect as determined by an unpaired, student’s t-test as follows: *, p <0.01; **, p < 0.001. Raw data for this figure can be viewed in Supplemental Table 3.

We also identified two alkaline-sensitive mutants. Loss of the bifunctional synthase PBP1a (*mrcA*) and the lytic transglycosylase MltG (*yceG*) impaired, but did not abolish, growth at pH 6.9 (Δ*mrcA*, 5% decrease in MDT; Δ*yceG*, 10% decrease in MDT) and pH 8.2 (Δ*mrcA*, 11% decrease in MDT; Δ*yceG*, 22% decrease in MDT). Both mutants exhibited wild type growth rates at pH 4.8 (Figure 1A, C). Individual deletions in the genes encoding for the six L,D-transpeptidases, five carboxypeptidases, and four amidases failed to confer any pH-dependent defects in MDT, consistent with either a limited role of these enzymes in exponential phase growth (13,14) or additional layers of redundancy (Figure 1-Figure Supplement1).

### aPBP activity ensures fitness across a wide pH range

Given their dramatic and opposing impact on MDT under acidic and alkaline conditions, we elected to focus further efforts on understanding the contribution of the bifunctional aPBPs PBP1a and PBP1b to growth across a range of pH conditions. An accumulating body of evidence suggests the aPBPs play overlapping, and potentially redundant, roles in cytoskeletal-independent PG synthesis during growth in standard culture conditions (i.e. nutrient rich, neutral pH growth media aerated at 37°C) (12,29). Indeed, PBP1a and PBP1b are a synthetic lethal pair in *E. coli* in these conditions (20).

Based on their disparate pH-dependent growth defects, we hypothesized that PBP1a and PBP1b are specialized synthases whose activity is optimized for growth under distinct pH environments. Consistent with this model, cells defective in PBP1a (Δ*mrcA*) and PBP1b (Δ*mrcB*) displayed defects in MDT at discrete, non-overlapping pH ranges. Between pH 5.1-5.9, loss of PBP1b resulted in a 5-22% reduction in MDT compared to wild type cells (p < 0.01), and growth was abolished at pH values below 4.8 (Figure 2A). The growth of the parental strain was not prevented until pH 4.2, over half a pH unit lower than the mutant (Figure 2B). At pH values at or above 6.2, MDT of this mutant was indistinguishable from wild type cells. Importantly, pre-conditioning the mutant in acidic media (pH 5.5) did not abrogate the growth rate defect (Figure 2B), indicating that steady-state pH—rather than pH shock—underlies the defect in MDT. In contrast to loss of PBP1b, the MDT of cells defective for PBP1a was equivalent to the parental strain during growth in acidic media (pH range 4.8-5.9), yet this mutant reproducibly exhibited between a 4-15% decrease in MDT from pH 6.2 to 8.2 (p < 0.05). The growth rates of mutant and wild type strains were not statistically distinguishable at pH 8.4 (Figure 2A). Loss of PBP1c (*pbpC*), a third aPBP with an unclear role in cell wall metabolism (30), did not result in a defect in MDT at any pH tested (Figure 2-Figure Supplement 1).

**Figure 2.**
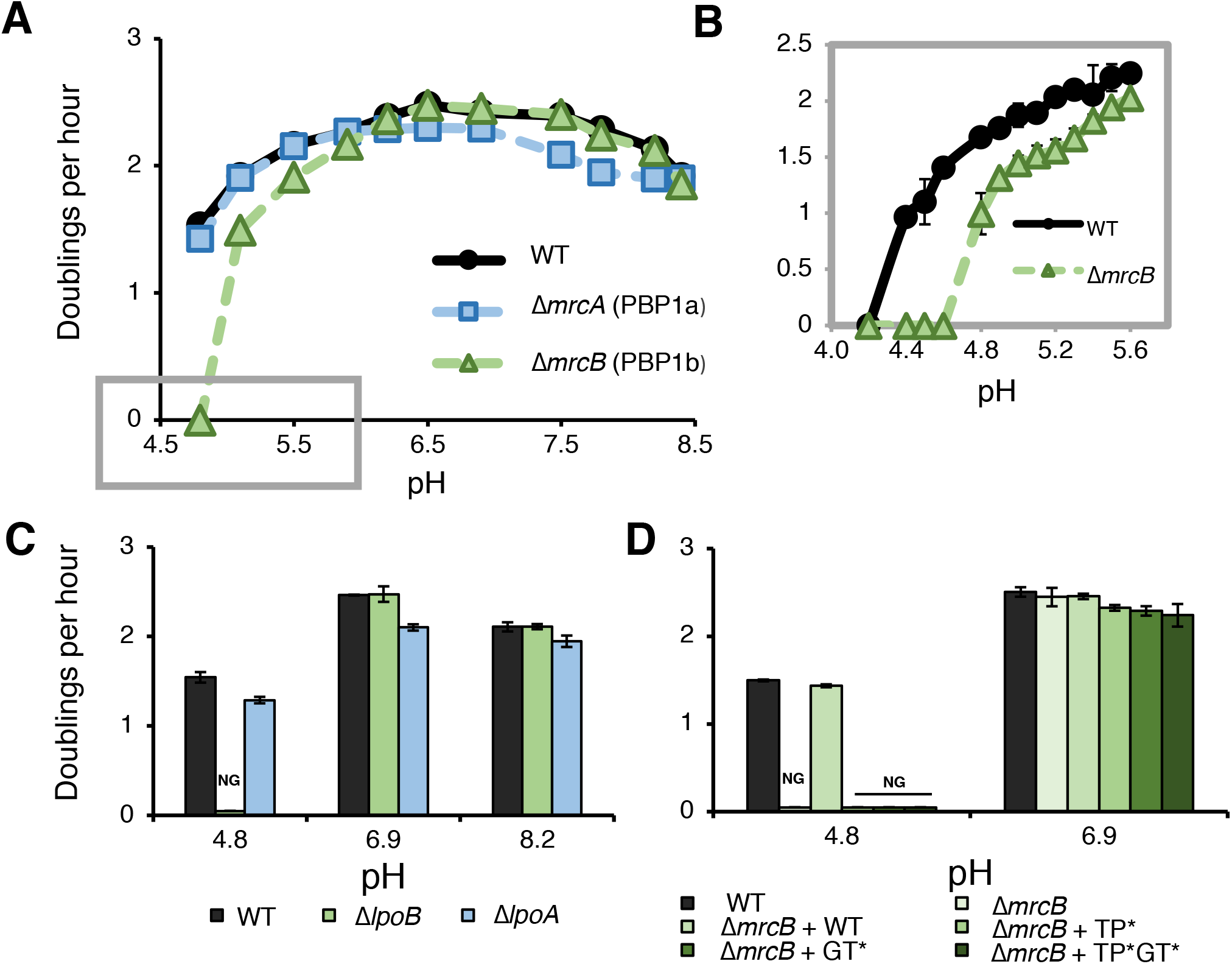
pH-dependent growth requires aPBP activity. A) Growth rate determination for Δ*mrcA* (PBP1a) and Δ*mrcB* (PBP1b) deletions compared to parental strain in LB media buffered from pH 4.8-8.4. Box drawn across x-axis indicates pH range assessed further in panel B. B) Growth rate of the wild-type strain and Δ*mrcB* in acidic conditions (pH 4.2-5.5) pre-cultured at pH 5.5. C) Growth rate analysis of *lpo* mutants cultured in buffered LB at pH 4.8, 6.9, or 8.0. D) Complementation analysis of PBP1b variants expressed from a plasmid and induced with 5 μM IPTG in the Δ*mrcB* background in buffered LB at pH 4.8 and 6.9. Markers represent average doublings per hour +/- standard deviation from three independent biological replicates. NG denotes “no growth” observed throughout the course of the experiment (20 hours).

We next sought to test whether aPBP transpeptidase and/or glycosyltransferase activity were required for fitness across pH conditions, as opposed to an indirect, structural role for these enzymes in the formation of cell wall synthesis complexes (31,32). To test this, we took advantage of two sets of mutants: 1) insertional deletions in *lpoA* and *lpoB*— genes encoding outer membrane lipoprotein cofactors required for activity, but not expression or stability, of PBP1a and PBP1b, respectively (33-36) and 2) point mutations that inactivate PBP1 b transpeptidase and/or glycosyltransferase activity but do not impact stability (37).

Implicating aPBP activity in growth across pH environments, loss of the cofactors LpoA and LpoB mimicked the pH-dependent growth defects of loss of the enzymes themselves. Analogous to cells defective for PBP1b, deletion of *lpoB* prevented growth at pH 4.8. Likewise, loss of PBP1a’s cofactor LpoA led to a growth rate defect at pH 6.9 and pH 8.2, although this mutant also exhibited a slight, but statistically significant, defect in MDT at pH 4.8 (Figure 2C). Complementation analysis of PBP1b variants at acidic pH further bolstered the conclusion that activity is required for pH-dependent growth. As expected, production of wild-type PBP1b *in trans* restored growth of the Δ*mrcB* mutant at pH 4.8; however, production of PBP1b variants rendering the transpeptidase (S510A, TP*), glycosyltransferase (E233Q, GT*), or both enzymatic activities inactive (TP*GT*) failed to complement growth (Figure 2D). It should be noted that consistent with observations that aPBP transpeptidase activity cannot be assayed *in vitro* in the absence of functional glycosyltransferase activity, the mutation in the glycosyltransferase active site (E233Q) previously has been shown to attenuate transpeptidase activity by 90% (35,38-40); thus, although our data demonstrate that transpeptidase activity is critical for pH-dependent growth, we cannot discern whether glycosyltransferase activity alone is required.

### aPBP activity promotes cell wall integrity across pH environments

Although these findings establish PBP1a and PBP1b activity as essential for optimal fitness across a wide pH range, it remained unclear whether these mutants’ pH-dependent defects in MDT in bulk culture were due to reduction in growth across the population (i.e. decreased rate of mass accumulation and cell expansion) or lysis of a fraction of cell in the population. To differentiate between these two mechanisms, we inoculated early exponential phase (OD_600_ ~ 0.05-0.1) wild type or mutant cells pre-grown at pH 6.9 on to agarose pads buffered to pH 4.5 or pH 8.0 and examined cells for incorporation of the dye propidium iodine (PI), which permeates cells with compromised membranes, by microscopy.

The pH-dependent growth defects were lytic in origin: extensive cell death was observed at one hour post-shift for PBP1b and PBP1a mutants that underwent acid (pH 6.9 to pH 4.5) or alkaline (pH 6.9 to pH 8.0) shock, respectively (Figure 3A). Time-lapse imaging of cells following pH shift shed light on lysis kinetics: upon pH downshift, Δ*mrcB* cells began to incorporate PI by 30 minutes (~15% cells labeled), with ~95-100% of cells labeled by two hours post-shift. Negligible cell death was observed for the parental strain or for cells defective for PBP1a during equivalent acid shock. Conversely, up to 15% of Δ*mrcA* cells underwent lysis an hour following alkaline shift (pH 6.9 to pH 8.0), comparable to the 10-15% growth defect observed in bulk culture at similar pH shifts (Figure 3B; Figure 2A). Δ*mrcB* and wild type cells exhibited minimal (< 5%) or negligible cell death, respectively, in response to alkaline shift. Interestingly, recovery of the *ΔmrcA* mutant was observed 90 minutes post-alkaline shock, suggesting cell damage may be sensed and initiate a feedback mechanism to restore cell wall integrity and fitness.

**Figure 3.**
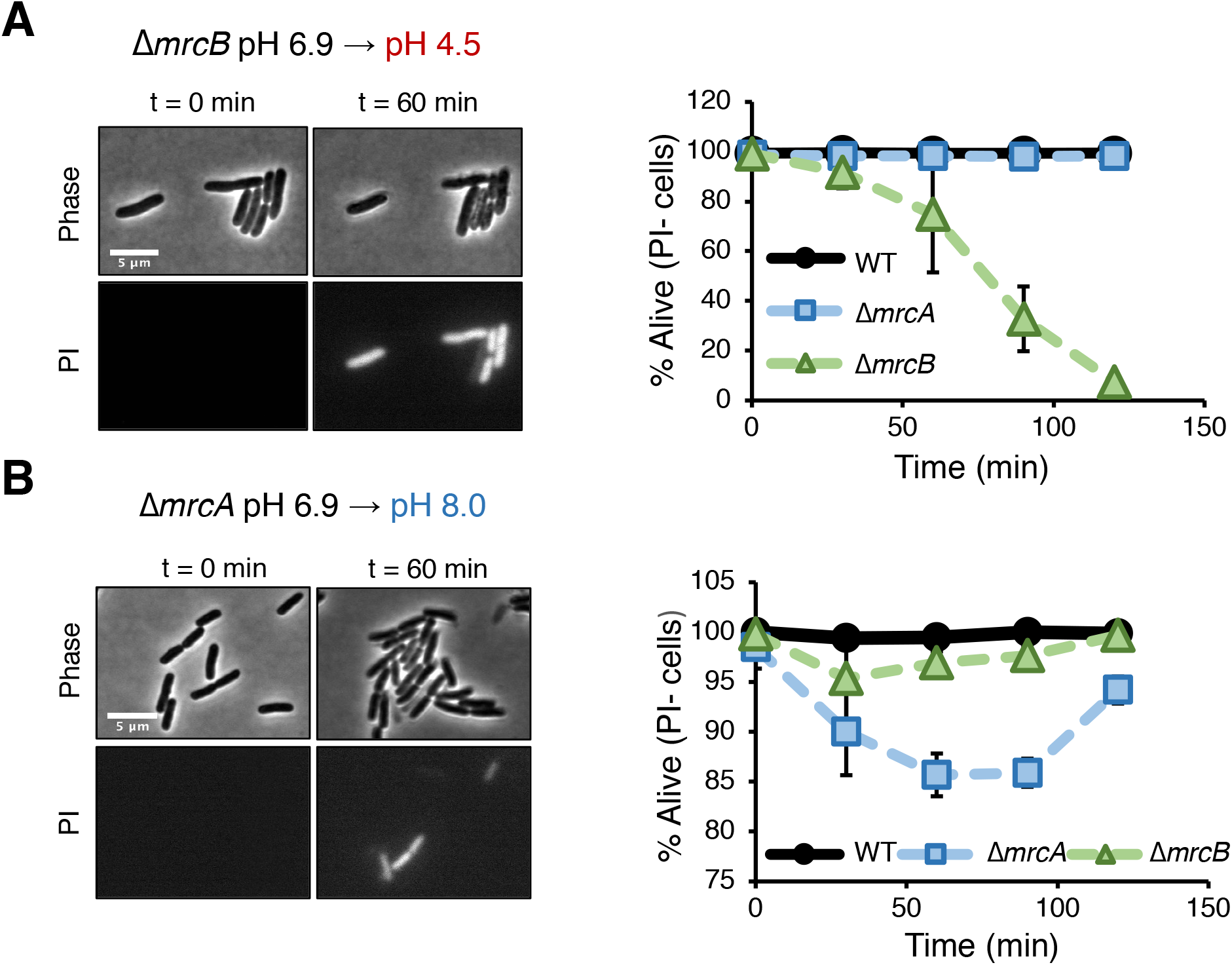
Mutants in aPBPs lyse upon exposure to non-permissive pH conditions. Micrographs depicting representative images of propidium iodine incorporation of *mrcA* (B, left) and Δ*mrcB* (A, left) mutants at t = 0 or 60 minutes post indicated pH shift. Scale bar represents 5 μm. B, Cell viability curves for wild-type, Δ*mrcA* (PBP1a), and Δ*mrcB* (PBP1 b) strains after acidic (A, right) or alkaline pH (B, right) shift as indicated. Cell death was determined by uniform staining of the cell boundaries with propidium iodine. Markers represent average percent viability +/- standard deviation of three biological replicates. Greater than 100 cells were analyzed for each strain at each time point per replicate (n = 3).

In addition to displaying differential lysis kinetics under their respective non-permissive pH conditions, PBP1a and PBP1b mutants also differed in apparent lysis mechanism. Time lapse imaging during acid shock revealed a high fraction of Δ*mrcB* cells lysed during division, often from a bulge emanating at the septum (Figure 4A, white arrows; Figure 4-Figure Supplement Movie 1). Scanning electron microscopy confirmed the bulges were coincident with the septum in this mutant (Figure 4C). To quantitate the lytic phenotype of the mutants, we categorized the lysis mechanism into three groups: septal bulge, non-septal bulge (including polar and peripheral bulging), and lysis not associated with visible bulging. Septal bulging was determined based on association of the bulge origin with the visible constriction site by phase contrast microscopy. Indeed, this analysis confirmed our observation: 55% of Δ*mrcB* mutants lysed at the septum following pH downshift with the remaining fraction associated with a non-septal bulge (17%) and no bulge (28%) (Figure 5D). In contrast, lysis of the Δ*mrcA* mutant during growth in alkaline pH was not associated with division, and instead, lysis of ~60% of the cells was coincident with a non-septal bulge, typically emanating from the periphery (Figure 4B, D; Figure 4-Figure Supplement Movie 2). These differences may reflect distinct localization preferences of the enzymes (32) or disparate weak regions in the PG across pH conditions.

**Figure 4.**
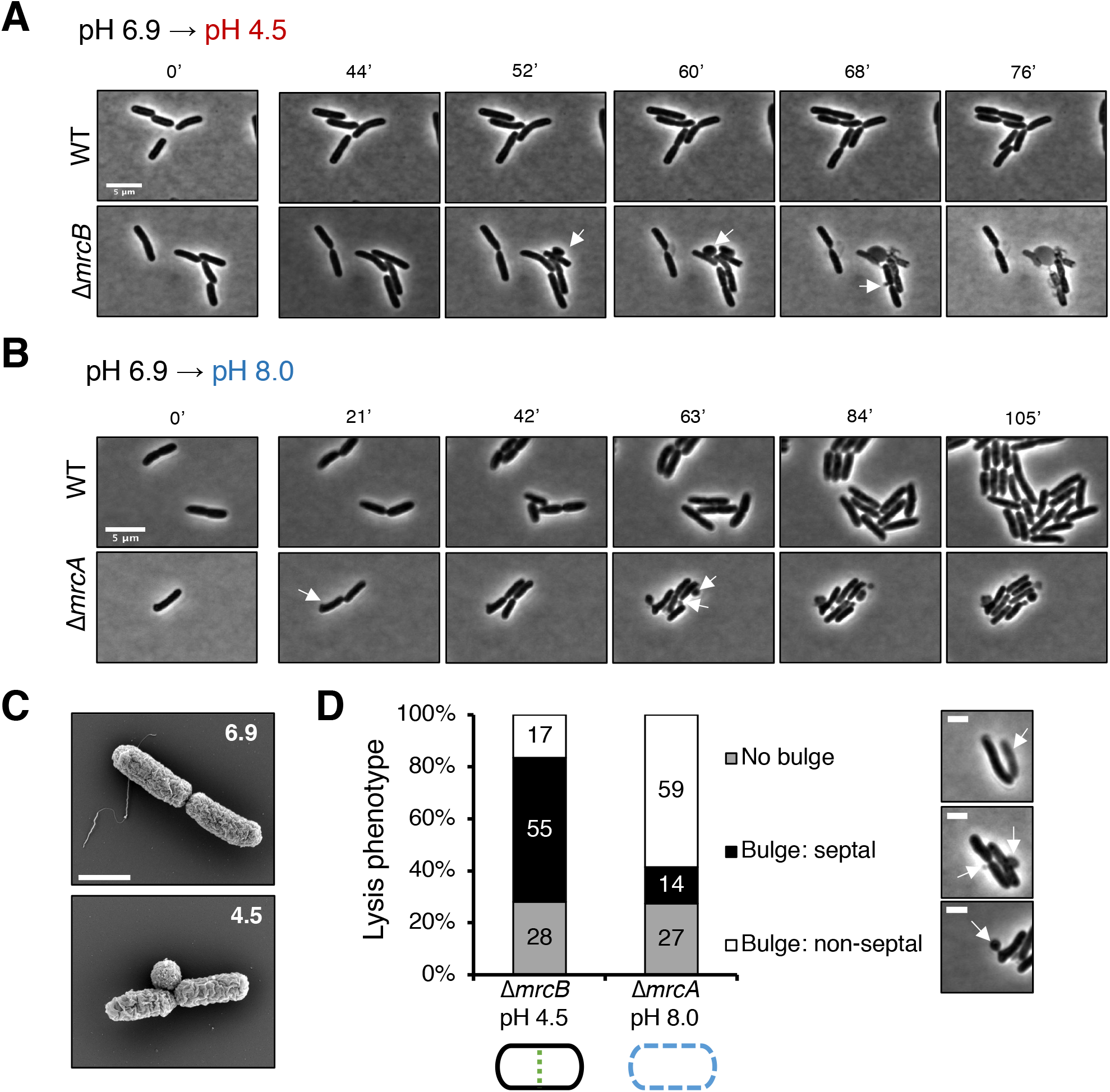
Distinct lytic phenotypes for cells defective for PBP1a and PBP1b upon pH shift. A-B) Representative phase contrast frames of time lapse imaging of Δ*mrcB* (PBP1b) and Δ*mrcA* (PBP1a) mutants upon acidic (A) or alkaline (B) pH shift, respectively, as compared to the parental strain. White arrows indicate membrane bulges. C) Representative scanning electron microscopy micrographs for Δ*mrcB* (PBP1 b) mutant shifted to either pH 6.9 or pH 4.5 for two hours prior to fixation. Scale bar represents 1 μm. D) Quantification of lysis phenotype between mutants. Lytic terminal phenotype was categorized into three groups: lysis via septal bulge, non-septal bulge, or no bulge. Determination of lytic phenotype was based on the frames preceding propidium iodine incorporation (time step = 3 minutes). Micrographs (top to bottom) depict representative images of no bulge, septal bulge, and non-septal bulge, respectively with arrows (scale bar = 2 μm). At least 50 cells across at least two independent biological replicates were assessed (Δ*mrcA, n* = 128; Δ*mrcB, n* = 278 cells). Bars are subdivided based on percent lytic phenotype in each mutant.

**Figure 5.**
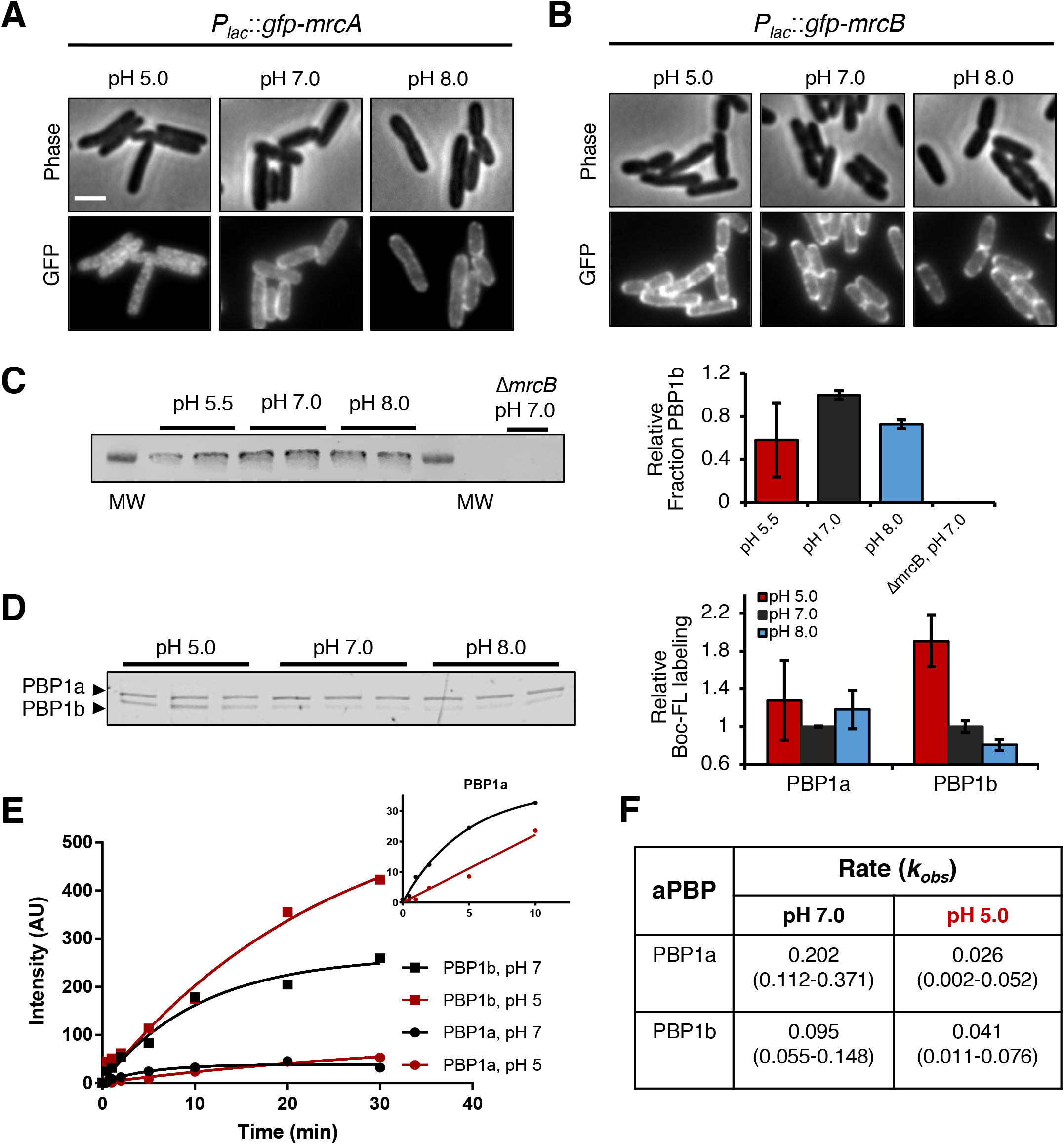
pH-dependence of aPBP localization, production and activity. A, B) Representative micrographs illustrating aPBP localization in strains Δ*mrcA P_lac_.gfp-mrcA* and Δ*mrcB P_lac::-gfp-mrcB_* grown in AB minimal media supplemented with 0.2% maltose and 250 μM IPTG at pH 5.0, 7.0 and 8.0. Scale bar indicates 2 μm. C) Western blot depicting two biological replicates of PBP1b production at pH 5.5, 7.0, and 8.0 with mean intensity +/- standard deviation (normalized to mean value at pH 7.0) shown to the right. The uncropped gel and the Ponceau staining for total protein levels can be viewed in Figure 5-Figure Supplement 1. D) Representative Bocillin-FL labeling of PBP1a and PBP1b from cell membranes incubated at pH 5.0, 7.0, and 8.0 for 15 minutes, each with three technical replicates shown. Quantification of PBP1a and PBP1b Bocillin-FL labeling as a function of pH is shown to the right. Bars represent the mean labeling (normalized to pH 7) +/- standard error of the mean from three separate membrane extractions, each including three technical replicates. E) Quantification of Bocillin-FL labeling to PBP1a and PBP1b at pH 5.0 and pH 7.0 across a time course (*t* = 0, 0.5, 1, 2, 5, 10, 30 min). Insert depicts early time point labeling of PBP1a. F) Apparent reaction rate (*k_obs_*) as a function of aPBP and pH calculated by nonlinear regression of data in E. Values in parentheses indicate 95% confidence intervals.

### aPBP substrate binding is pH-dependent

Although our data support a model in which aPBP activity is differentially required for fitness across pH environments, the mechanistic basis for pH specialization remained unknown. To interrogate this, we compared the production, localization, and substrate binding of PBP1a and PBP1b as a function of pH. Consistent with previous a proteomic mass spectrometry dataset observing limited differences in abundance of either aPBP across pH conditions (41), bulk protein levels of PBP1b were not significantly influenced by pH (Figure 5C). Likewise, similar to previous reports (32,42), green fluorescent protein fusions to PBP1a and PBP1b expressed at an ectopic site exhibited discrete localization profiles: PBP1a localized solely to the cell periphery, while PBP1b was present at both the periphery and the septum. However, few localization differences were observed across pH conditions with the exception of GFP-PBP1a, which appeared to adopt an irregular clustered distribution throughout the cell body at pH 5.0 (Figure 5A, B).

To probe the effect of pH on substrate binding, we compared the ability of aPBPs to bind substrate analog Bocillin-FL (Boc-FL), a fluorescent penicillin derivative. Analogous to β-lactam antibiotics, Boc-FL labeling requires an accessible catalytic serine in the transpeptidase active site for acylation, and thus labeling has been used to assay the efficiency of a key step of the transpeptidation reaction (23,43). Although not a direct readout of enzymatic activity, Boc-FL labeling offers numerous advantages over *in vitro* synthase assays. Notably, Boc-FL labeling can measure activity of multiple PBPs simultaneously and can be performed in live cells or in membrane extracts, thus ensuring more physiologically relevant labeling conditions, which may have occluded observation of pH-dependent differences in PBP1b activity in previous *in vitro* studies (35). To ensure invariable abundance of all PBPs in our experiments, we extracted membranes from wild-type cells grown at neutral pH and incubated membranes with Boc-FL in buffer to a final pH of 5.0, 7.0, or 8.0.

PBP1b binding to Boc-FL was strikingly pH-dependent, suggesting that at least one aspect of its transpeptidase reaction is enhanced in acidic environments. After a 15-minute incubation with Boc-FL, PBP1b labeling was elevated by 200% at pH 5.0 and decreased by 20% at pH 8.0 compared to neutral pH (Figure 5D; p < 0.05), consistent with previous results using radiolabeled penicillin (44). Further examination of PBP1b binding kinetics at pH 5.0 and pH 7.0 revealed that while reaction rate was independent of pH, the saturation point varied (Figure 5E, F), suggesting an increase in the active fraction of molecules at acidic pH (12). In single time point or kinetic assays, PBP1a binding to Boc-FL saturated at similar levels across pH values. However, rate of Boc-FL binding for the synthase was reduced approximately 10-fold at acidic pH (Figure 5E, F), indicating that PBP1a has reduced activity in this condition.

### pH-dependent PBP1b activity alters intrinsic resistance to PBP2 and PBP3 specific β-lactam antibiotics

PBP1b activity has previously been implicated in intrinsic resistance to β-lactam antibiotics, specifically to compounds that target the elongation specific transpeptidase PBP2 and the division specific transpeptidase PBP3. Strains defective for PBP1b or lipoprotein activator LpoB show enhanced susceptibility to PBP2/PBP3-specific compounds through rapid lysis (42,45,46), and elevated PBP1b activity was recently shown to protect against PBP2 specific antibiotic mecillinam (15).

Given our finding that low pH enhances PBP1b activity and previous reports of β-lactam tolerance in acidic media (47), we predicted that growth in acidified media may decrease susceptibility to antibiotics that specifically inactivate PBP2 and PBP3, and that the observed resistance should require PBP1b activity. If true, condition-dependent intrinsic resistance may have important implications for treatment of *E. coli* infections in host niches with variable pH (26,27). To test this model, we exposed our wild type strain (MG1655) to a panel of β-lactam antibiotics and quantified the minimum inhibitory concentration (MIC) across the physiological pH range of 4.5-8.0.

In support of our hypothesis, we observed a 2 to 64-fold increase in MIC to compounds that selectively inactivate PBP2 and PBP3 (48) at pH values < 6.0 (Figure 6A). Acidic pH appeared to confer a protective effect on the elongation and division machinery: cells cultured in low pH media retained near-normal morphology in the presence of concentrations of the compounds that led to either filamentation (cephalexin, CEX) or cell rounding (mecillinam, MEC) at pH 7.0 (Figure 6B, C). In contrast, susceptibility to a non-specific β-lactam (ampicillin, AMP), an aPBP-targeting compound (cefsulodin, CFS) (48), or a protein synthesis inhibitor (chloramphenicol, CH) was not strongly pH-dependent. We additionally confirmed that low pH-dependent resistance could not be attributed to stability of the antibiotics, loss of the proton motive force, or β-lactamase production (Figure 6-Figure Supplement 1, 2). Changes in outer membrane permeability, another common mechanism for β-lactam resistance, would be expected to affect all β-lactams equally and thus is unlikely to underlie pH-dependent resistance. Importantly, resistance was not limited to our laboratory strain, as uropathogenic *E. coli* isolate UTI89 (49) exhibited a comparable pH-dependent change in MIC to both cephalexin (CEX) and mecillinam (MEC) at low pH during growth in both broth culture and in urine (Figure 6D).

**Figure 6.**
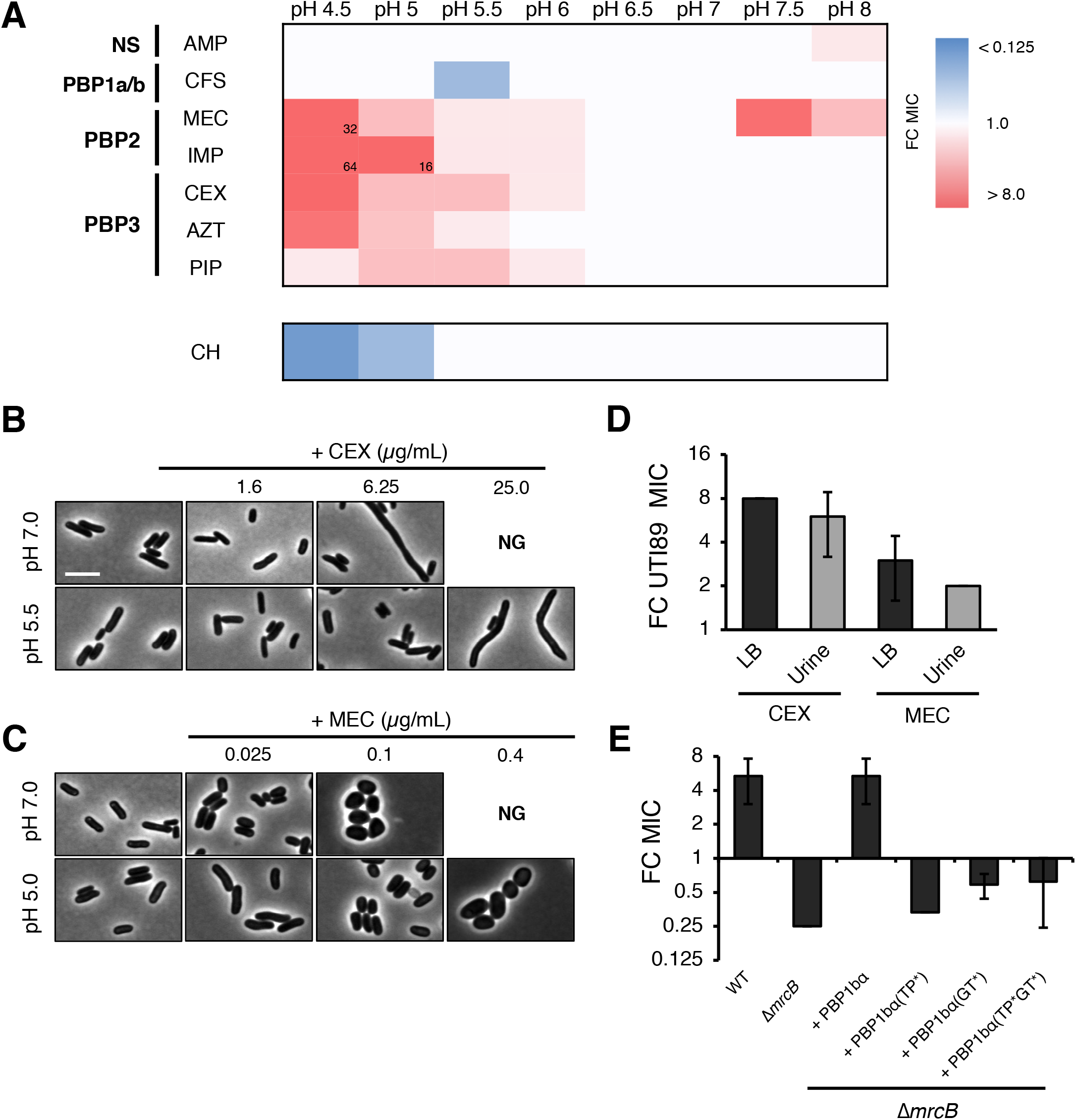
Intrinsic resistance to PBP2 and PBP3-targeting β-lactams at low pH. A) Heat map summarizing fold change in minimum inhibitory concentrations (MIC) of antibiotics for strain MG1655 cultured in LB (pH 4.5-8.0) after 20 hours. Cells in heat map are colored based on median fold change (FC) in MIC at indicated pH compared to pH 7.0 from at least three biological replicates. Raw MIC values can be viewed in Supplemental Table 3. Abbreviations for antibiotic names are as follows: AMP, ampicillin; CFS, cefsulodin; IMP, imipenem; MEC, mecillinam; CEX, cephalexin; AZT, aztreonam; PIP, piperacillin; CH, chloramphenicol. Fold change values of > 8 are indicated in the lower right-hand corner of the cell. B-C) Representative micrographs of terminal phenotypes of cells treated with PBP3 inhibitor cephalexin (B) or PBP2 inhibitor mecillinam (C) cultured at pH 7.0, pH 5.5 or pH 5.0. Scale bar indicates 3 μm. NG denotes “no growth” observed at the indicated concentration of antibiotic. D) Fold change in minimum inhibitory concentration of *E. coli* strain UTI89 to cephalexin (CEX) and mecillinam (MEC) grown at pH 5.0 compared to pH 7.0 in broth culture and in urine. Raw MIC values can be viewed in Supplemental Table 4. E) Fold change in minimum inhibitory concentration to cephalexin (CEX) for indicated strains grown at pH 5.5 compared to pH 7.0. Cells producing PBP1b variants were grown in the presence of 10 μM IPTG. Raw MIC values can be viewed in Supplemental Table 5. Bars represent average fold change in minimum inhibitory concentration +/- standard deviation across at least three biological replicates.

Genetic analysis suggests that PBP1b activity is specifically required for pH-dependent resistance to β-lactam antibiotics. Strains defective in non-essential transpeptidases, including aPBPs PBP1a and PBP1c and L,D-transpeptidases YcbB and YnhG, had no impact on resistance to the PBP3 inhibitor cephalexin (CEX). In contrast, loss of PBP1b abolished resistance at pH 5.5 and in fact, slightly increased susceptibility (Figure 6D). Resistance to other PBP2 and PBP3 targeting compounds was also eliminated or significantly reduced in cells defective for PBP1b (Figure 6-Figure Supplement 3C). Moreover, PBP1 b activity was required for resistance: loss of the enzyme’s cognate outer membrane lipoprotein LpoB or inactivation of its catalytic activity eliminated CEX resistance (Figure 6E, Figure 6-Figure Supplement 3B). Importantly, a mutant with a comparable growth rate defect at pH 5.5 (Δ*mrcB* 1.91 +/-0.02 doublings per hour; Δ*tolA* 1.40 +/-0.04 doublings per hour) still displayed the same fold change in resistance to CEX at pH 5.5 as the parental strain. Likewise, a mutant in PBP5 (Δ*dacA*) with increased sensitivity to CEX even at neutral pH, similar to Δ*mrcB* (45,46), also retained the resistance phenotype in acidic growth conditions (50) (Figure 6-Figure Supplement 3D).

## DISCUSSION

### Specialization role for aPBPs in cell wall integrity across environmental conditions

By varying a single growth parameter—extracellular pH—we were able to uncover specialized roles for a subset of nonessential cell wall synthases and autolysins in *E. coli* that had previously been classified as redundant. Of the pH specialist enzymes identified, we chose to focus on the bifunctional synthases PBP1a and PBP1b, which we find are required for cell wall integrity in distinct pH environments in part due to pH-dependent changes in substrate binding (Figure 5D-F). Both PBP1a and PBP1b exhibit reduced capacity for substrate binding outside of their preferred pH range, albeit through distinct mechanisms (e.g. change in rate of binding for PBP1a vs. change in binding saturation for PBP1b). In effect, they are unable to fully compensate for loss of the other during growth in particular pH environments, leading to cell lysis and reduced fitness (Figure 7). Apart from activity, the spatial preferences of the enzymes are likely to contribute to specialization as well: for example, PBP1a peripheral localization may lead to unfilled gaps in mid-cell PG and thus contribute to the septal lysis phenotype of the *DmrcB* mutant (Figure 5). Importantly, as few differences in peptidoglycan composition are observed in *E. coli* cells grown in pH 7.5 and pH 5.0 media, it seems likely that the differential requirement for the aPBPs reflects a change in pH-dependent changes in enzymatic activity rather than altered cell wall structure (23).

**Figure 7.**
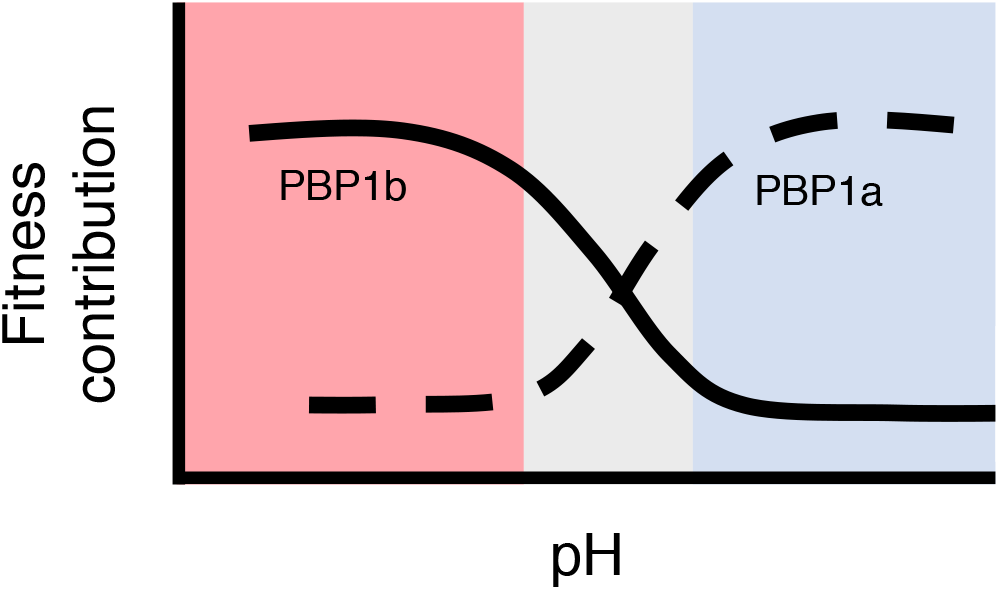
Model depicting contribution of aPBPs to fitness across pH environments.

Interestingly, apart from displaying depressed capacity for substrate binding in alkaline conditions, PBP1 b also displays increased affinity for substrate in acidic conditions, where our data support it is the predominant aPBP. Furthering bolstering the enhanced activity of the enzyme in this environment, we found PBP1b mediates low pH-dependent resistance to PBP2 and PBP3-targeting β-lactam antibiotics (Figure 6E). Although these findings appear in conflict with biochemical data demonstrating PBP1b glycosyltransferase activity is reduced under acidic conditions *in vitro* (35), we reason that this discrepancy may indicate that pH-dependent changes to PBP1b activity may be mediated through a regulator rather than through a direct effect on enzymatic activity itself. Although the outer membrane activator LpoB is an attractive candidate, it is unlikely the sole activator, as it fails to significantly enhance PBP1b activity in acidic conditions (35). Other candidate activators include the terminal cell division protein FtsN, which interacts individually or in combination with LpoB to stimulate PBP1b glycosyltransferase activity *in vitro* (31,51), and the pH specialist autolysins we identified here (Figure 1B, C), several of which have been recently shown to activate PBP1b *in vivo*, potentially through generating gaps in the PG matrix (12,15,52). Future work will be aimed at determining the contribution of each of these factors to PBP1b activity across pH conditions and assessing whether pH-dependent PBP1a activity is regulated by intrinsic or extrinsic factors.

Although PBP1a and PBP1b share an essential role in maintenance of cell wall integrity under standard culture conditions (20), there had been previous hints these enzymes were not completely interchangeable. Beyond possessing distinct interaction partners and subcellular localization profiles (31,32,53), mutants display differential susceptibility to antibiotic treatment, osmotic shifts, and mechanical stress (29,42,54). We anticipate further study of these synthases—as with other “redundant” cell wall autolysins—under nonstandard culture conditions will continue to reveal unique roles for these enzymes in cell wall biogenesis.

### Plasticity in cell wall metabolism potentiates intrinsic resistance to cell wall active antibiotics

Analogous to alternative PBP usage in methicillin resistant *Staphylococcus aureus* (55,56), we find a key consequence of environment-driven plasticity in *E. coli* cell wall metabolism is intrinsic resistance to β-lactam antibiotics with a narrow target specificity. The most commonly prescribed class antibiotics, β-lactam antibiotics are frequently used to treat *E. coli* infections, including in treatment of urinary tract infections. As the urine pH varies between and within individuals (27), it is possible that pH-dependent resistance may contribute to treatment failure of particular therapeutics in the clinic. Even in circumstances in which pH-dependent resistance to certain compounds appears mild (2 to 4-fold), incomplete resistance and tolerance may elevate the mutation rate, thereby accelerating emergence of complete resistance (57).

Although intrinsic resistance is a particularly concerning consequence of plasticity in cell wall synthesis, the mechanistic underpinnings for the role of PBP1b in pH-dependent resistance remain unclear. We imagine three models that may account for this finding: 1) at low pH PBP1b can partially substitute for PBP2 and PBP3 activity in elongation and division, respectively, 2) increased flux of PG precursors through PBP1b and away from PBP2 and PBP3 dampens the lethal effects of the futile cycle (15,58), or 3) PBP1b recognizes and repairs damaged PG matrix as a result of β-lactam activity, thus serving a quality control function. In light of our observation that cellular dimensions are maintained at higher concentrations of the compounds when the cells are cultured in acidic media (Figure 6B, C), we favor a model in which PBP1b partially replaces the monofunctional PBP2 or PBP3 in the Rod system and divisome, respectively, similar to what has previously been observed for bPBPs PBP2b and PBP3 in the *Bacillus subtilis* divisome (59). Decreased reliance on bPBP activity would be expected to render cells less sensitive to PBP2 and PBP3 specific antibiotics (60). It is unclear how proper morphology would be preserved if PBP1b-mediated resistance functioned through dampening the futile cycle (model 2) or through repairing damaged cell wall material (model 3). Although further work is necessary to discern which, if any, of these models underlie this phenotype, clarification of PBP1b’s role in resistance—especially across environmental conditions—promises to shed light on cell wall biogenesis and inform the design and use of novel antibiotic therapies.

### Apparent redundancy ensures fitness across environmental conditions

Although clearly of consequence for antibiotic resistance, environmental specialization of cell wall enzymes likely functions more broadly as a key adaptation allowing *E. coli* to thrive across an unusually wide pH range (pH ~4-9) and even tolerate extreme pH shocks, such as during transient exposure to gastric acid (pH ~2) (61). This physiological adaptation to preserve essential processes in the periplasm likely works in concert with the organism’s ability to modify extracellular pH through the export of acidic and alkaline substrates (62,63). In this context, pH-specialist cell wall enzymes may function in part to maintain cell wall integrity until the extracellular media reaches a growth-permissive pH.

Among other pH tolerant organisms, strategies employed to expand growth across wide pH ranges are likely to vary—even between closely related species. Instead of differential aPBP activity underling pH tolerance, *S. enterica* serovar Typhimurium encodes a PBP3 paralog, termed PBP3_SAL_, that is active during septation during growth in acidic environments, including in the organism’s intracellular lifecycle (24). As PBP3_SAL_ is restricted to *Salmonella, Enterobacter*, and *Citrobacter* spp., alternative mechanisms to cope with changing pH environments must exist. Elucidating the requirements for pH-dependent growth in organisms outside the *Enterobacteriaceae* will shed light on whether aPBPs, which are broadly conserved across bacteria (33), play a central role in the process.

Apart from pH, we anticipate enzyme specialists exist across environmental conditions; redundancy in cell wall enzymes is present throughout bacterial species, even among those that only grow at a narrow pH range (16). *B. subtilis*, for example, encodes 16 PBPs alone, yet its growth is restricted to pH 6.0-pH 9.0 (64). Ionic strength, osmolality, and temperature are among the properties that vary across the habitats bacteria occupy and may modulate the chemical and physical properties of the periplasm and the extracytoplasmic space of Gram-positive bacteria. In support of this, PBP2 from *Caulobacter crescentus* displays differential localization patterns as a function of extracellular osmolality (65), and the lytic transglycosylase MltA from *E. coli* has ~10 times greater activity at 30°C than at 37°C *in vitro* (25,66).

Beyond cell wall enzymes, we expect environmental specialization will underlie the high levels of redundancy in other periplasmic protein classes, including sugar transporters, efflux pump adapter proteins, and chaperones (67-69). Nevertheless, improved understanding of the contribution of many enzymes to bacterial fitness in the wild demands a departure from standard growth conditions.

## MATERIALS AND METHODS

### Bacterial strains, plasmids, and growth conditions

Unless otherwise indicated, all chemicals, media components, and antibiotics were purchased from Sigma Aldrich (St. Louis, MO). Bacterial strains and plasmids used in this study are listed in Supplemental Table 1 and Supplemental Table 2, respectively. All *E. coli* strains are derivatives of MG1655, and all deletion alleles were originally provided by the Coli Genetic Stock (70). Unless otherwise indicated, strains were grown in lysogeny broth (LB) media (1% tryptone, 1% NaCl, 0.5% yeast extract) supplemented with 1:10 MMT buffer (1:2:2 molar ratio of D,L-malic acid, MES, and Tris base) to vary media pH values between pH 4-9. Uropathogenic *E. coli* strain UTI89 was also cultured in urine provided by a healthy donor and supplemented with MMT buffer to fix the pH. When selection was necessary, cultures were supplemented with 50 μg/mL kanamycin (Kan) and 100 μg/mL ampicillin (Amp). Cells were grown at 37°C either in 96-well microtiter plates shaking at 567 cpm or in glass culture tubes shaking at 200 rpm for aeration.

### Growth rate measurements

Strains were grown from single colonies in glass culture tubes in LB + MMT buffer (pH 6.9) to mid-log phase (OD_600_ ~0.2-0.6), pelleted, and re-suspended to an OD_600_ 1.0 (~1 x 10^9^ CFU/mL). Cells were diluted and inoculated into fresh LB + MMT buffer at various pH values in 96-well plates (150 μl final volume) at 1 x 10^3^ CFU/mL. Plates were grown at 37°C shaking for 20 hours in a BioTek Eon microtiter plate reader, measuring the OD_600_ of each well every ten minutes. Doublings per hour was calculated by least squares fitting of early exponential growth (OD_600_ 0.005-0.1).

### Microscopy and time lapse imaging

For time lapse imaging experiments, cells were grown from a single colony in LB + MMT buffer (pH 6.9) to early exponential phase (OD_600_ ~0.05-0.1) then mounted onto 1.0% agarose pads at pH 4.5, 6.9, or 8.0. Where indicated, propidium iodine was added to the agarose pad at a final concentration of 1.5 μM. Cells were allowed to dry on pads 10 minutes prior to imaging. All phase contrast and fluorescence images were acquired on a Nikon Ti-E inverted microscope (Nikon Instruments, Inc.) equipped with a 100X Plan N (N.A. = 1.45) Ph3 objective, X-Cite 120 LED light source (Lumen Dynamics), and an OrcaERG CCD camera (Hammamatsu Photonics, Bridgewater, N.J.). Filter sets were purchased from Chroma Technology Corporation. The objective was pre-heated to 37°C using an objective heater. Image capture and analysis was performed using Nikon Elements Advanced Research software. Cell death quantification was determined by cells uniformly stained with propidium iodine, and terminal lytic phenotype of cells was determined by assessment of the frames immediately preceding propidium iodine incorporation.

For aPBP localization studies, cells were grown in AB minimal media supplemented with 0.2% maltose and 250 μM IPTG overnight then sub-cultured into fresh media the following morning. Cells were grown to OD_600_ 0.1-0.2 at 37°C then fixed by adding 20 μL of 1M NaPO4, pH 7.4, and 100 μL of fixative (fixative = 1 mL 16% paraformaldehyde + 6.25 μL 8% glutaraldehyde). Samples were incubated at room temperature for 15 min, then on ice for 30 min. Fixed cells were pelleted, washed three times in 1 mL 1X PBS, pH 7.4, then resuspended in GTE buffer (glucose-tris-EDTA) and stored at 4°C.

### Scanning electron microscopy

Wild type and Δ*mrcB* cells were grown to mid-exponential phase in MMT buffered pH 6.9 LB media and back-diluted to an OD_600_ = 0.1 into either pH 6.9 or 4.5 media. Cells were allowed to grow for an additional hour, fixed as described above, and applied to polylysine coated coverslips. Post fixation, samples were rinsed in PBS 3 times for 10 minutes each followed by a secondary fixation in 1% OsO4 in PBS for 60 minutes in the dark. The coverslips were then rinsed 3 times in ultrapure water for 10 minutes each and dehydrated in a graded ethanol series (50%, 70%, 90%, 100% x2) for 10 minutes each step. Once dehydrated, coverslips were then loaded into a critical point drier (Leica EM CPD 300, Vienna, Austria) which was set to perform 12 CO2 exchanges at the slowest speed. Once dried, coverslips were then mounted on aluminum stubs with carbon adhesive tabs and sputter coated with 6nm of iridium (Leica ACE 600, Vienna, Austria). After coating, the samples were then loaded into a FE-SEM (Zeiss Merlin, Oberkochen, Germany) imaged at 3 KeV with a probe current of 178 pA using the Everhart Thornley secondary electron detector.

### Bocillin-FL labeling of membrane extract

Cells were grown from a single colony in a 25 mL flask overnight, sub-cultured 1:200 into 1 L of LB media the next morning and grown to an OD_600_ 1.4-1.6. To extract the cell membrane, cells were pelleted at 4,400 x *g* for 10 minutes at 4 °C, resuspended in buffer (10 mM potassium phosphate + 140 mM NaCl, pH 7.0), and then sonicated. Cell lysate was collected, and total protein concentration was measured by Bradford assay after centrifugation at 9,100 x *g* for 10 min at 4 °C. 10 mg/mL of crude membrane preparation was incubated in the presence of buffer (20 mM potassium phosphate + 140 mM NaCl at varying pH values) and 0.1 mM Bocillin-FL (Thermo Fisher Scientific, Waltham, MA) for indicated time interval at 37 °C in the dark, and then the reaction was quenched upon addition of Laemmli buffer to 1x final concentration. Samples were boiled for 10 minutes, then 20 μL was separated on a 12% SDS-PAGE gel in the dark. Gels were imaged on a BioRad XR+ gel imager (Alexa488 setting), and background subtraction and band quantification was performed in ImageJ. For time course experiments, apparent reaction rate (*k_obs_*) was calculated in GraphPad Prism by nonlinear fitting of one phase early exponential decay with the constraint of Y_0_ = 0.

### SDS-PAGE and immunoblotting

Strains were grown from a single colony in LB prepared at pH 5.5, 7, or 8 to mid-log phase (OD_600_ ~ 0.2-0.6), back-diluted, and grown to an OD_600_ ~ 0.4 to achieve balanced growth. An aliquot of each culture was sampled to ensure the pH of the culture remained unchanged from the starting pH value. Samples were pelleted, re-suspended in 2x Laemmli buffer to an OD_600_ ~ 20, and boiled for ten minutes. Samples (10 μl) were separated on 12% SDS-PAGE gels by standard electrophoresis, transferred to nitrocellulose membranes, and probed with PBP1b antisera. PBP1b antisera (rabbit) was used at a 1:1000 dilution, and an HRP-conjugated secondary antibody (goat α-rabbit) was used at a 1:2000 dilution. Blots were imaged on a LiCor Odyssey imager. Quantitation of relative PBP1b protein levels were determined in ImageJ and normalized to Ponceau staining as a loading control.

### Antibiotic susceptibility testing

For determination of minimum inhibitory concentrations, cells were grown from a single colony in LB media at the indicated pH to mid-exponential phase (OD_600_ ~ 0.2-0.6) at 37°C with aeration and then inoculated at 1 x 10^5^ CFU/mL into LB media of the same pH in sterile 96-well plates with a range of two-fold dilutions of the indicated antimicrobial agent (final volume, 150 μL). Plates were incubated at 37°C shaking for 20 hours before determination of the well with the lowest concentration of the antibiotic that had prevented growth by visual inspection.

### Antibiotic stability testing

Antibiotics were incubated for 20 hours in LB media at pH 4.5, 7.0, or 8.0 and then diluted into 96-well plates containing LB media (pH 7) and 1 x 10^5^ CFU/mL MG1655. Plates were then incubated at 37°C shaking for 20 hours before determination of the compound’s minimum inhibitory concentration.

### Terminal phenotype assessment

Cells from minimum inhibitory concentration assays were spotted (5 μL) onto 1.0% agarose pads 20 hours post-treatment and imaged by phase contrast microscopy to track cell morphology in response to antibiotic treatment across pH values. Growth rate was monitored by OD_600_ in the BioTek Eon plate reader to confirm all cells examined were in the same growth phase and at approximately the same optical density prior to imaging.

### Quantification and statistical analysis

A minimum of three biological replicates were performed for each experimental condition. Data are expressed as means +/- standard deviation or standard error of the mean. Where appropriate, a student’s two-tailed unpaired t-test was applied with a significance threshold set to p < 0.05 or < 0.01 (Figure 1). Asterisks indicate significance difference as described in corresponding figure legend.

## ACKNOWLEDGEMENTS

We thank Waldemar Vollmer (plasmids, PBP1 b anti-serum) and Tom Bernhardt (strains) for kind gifts critical for completion of this work. We thank Pam Brown and Michelle Williams for advice on the Bocillin-FL labeling procedure, the Goldberg lab for use of their gel imager, and Abbygail Iken for technical assistance with the UTI89 MIC assays. We gratefully appreciate sample preparation and electron microscopic imaging assistance from Matthew Joens, Daniel Geanon, Greg Strout and Dr. James Fitzpatrick from the Washington University Center for Cellular Imaging which is supported by Washington University School of Medicine, The Children’s Discovery Institute of Washington University and St. Louis Children’s Hospital (CDI-CORE-2015-505) and the Foundation for Barnes-Jewish Hospital (3770). We are indebted to members of the Levin and Zaher labs for fruitful discussions on technical and philosophical matters related to this this research, as well as Corey Westfall and Joseph Merriman for critical reading of this manuscript. This work was supported by NIH grants GM64671 and GM127331 to P.A.L. and NSF Graduate Research Fellowship DGE-1745038 to E.A.M.

---

**Figure 1- Figure Supplement 1.**
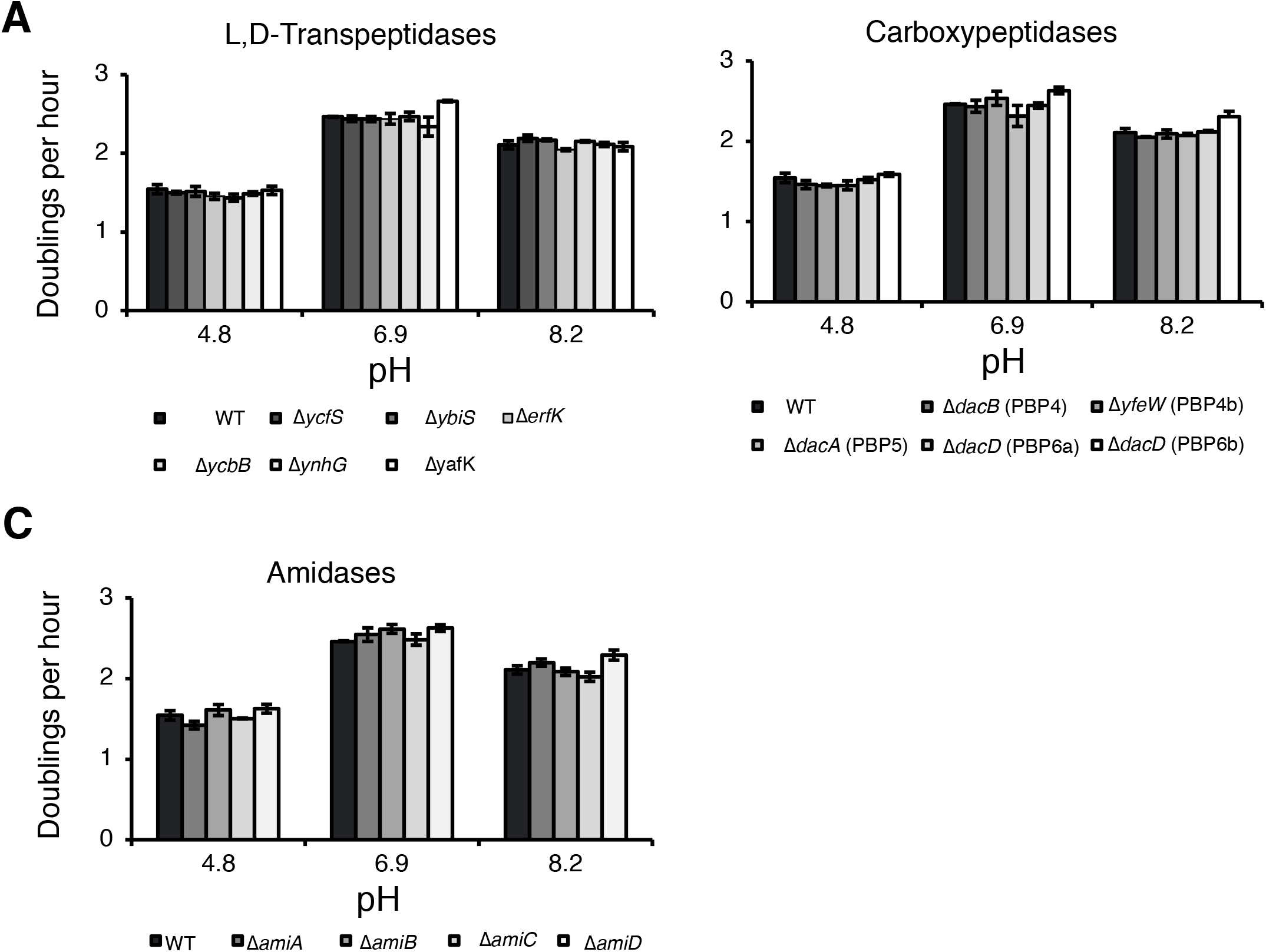
Growth rate of cell wall mutants across pH environments. Insertional deletions in non-essential L,D-transpeptidases (A), carboxypeptidases (B), and amidases (C) were screened for growth defects compared to the parental strain in LB media buffered to pH 4.8, 6.9, and 8.2. Bars depict average doublings per hour +/- standard deviation of three independent biological replicates. Raw data for this figure can be viewed in Supplemental Table 3.

**Figure 2- Figure Supplement 1.**
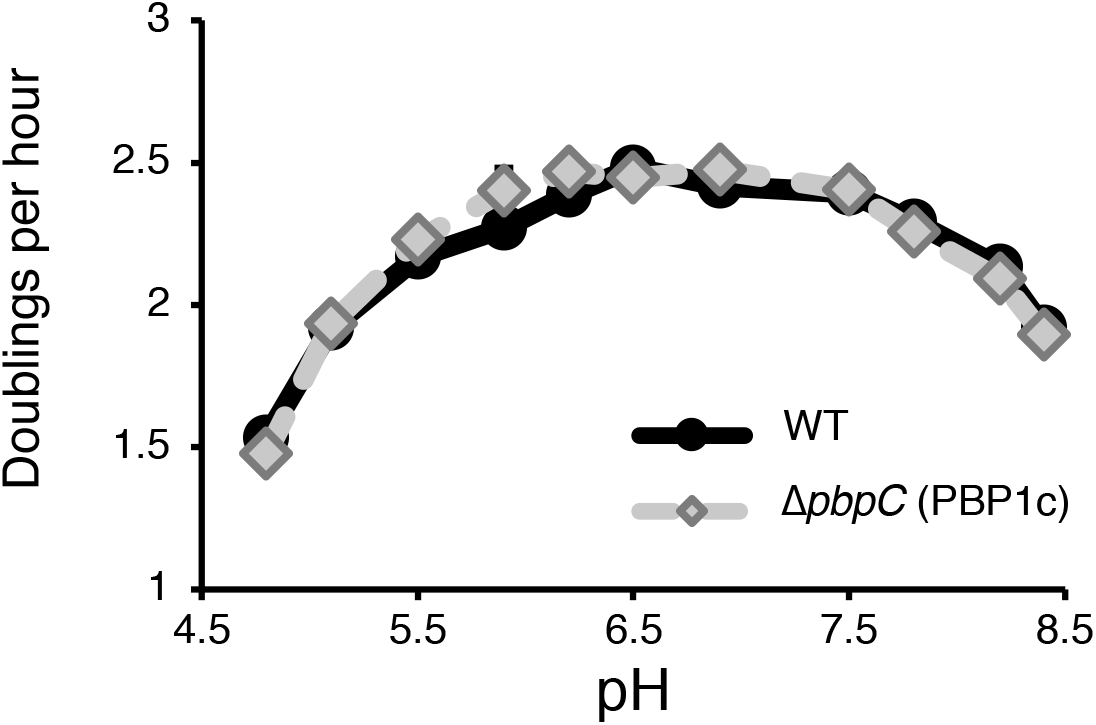
Growth rate analysis of strain defective for PBP1c enzyme. Growth rate determination for Δ*pbpC* (PBP1c) compared to parental strain in LB media buffered from pH 4.8-8.4. Markers depict average doublings per hour +/- standard deviation of three independent biological replicates.

**Figure 5- Figure Supplement 1.**
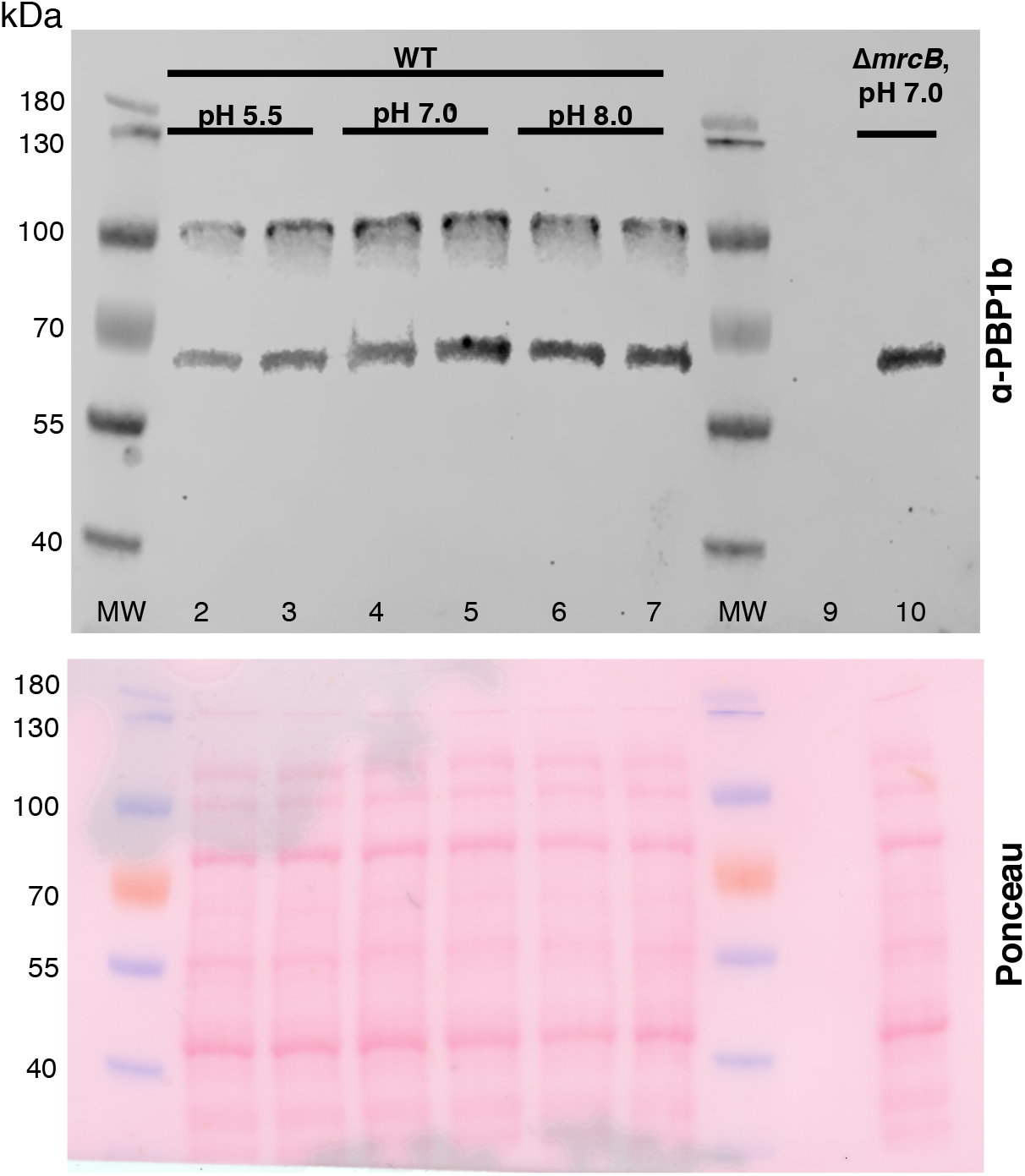
Production and localization of PBP1b as a function of pH. Full immunoblot and corresponding Ponceau stain for blot shown in Fig. 5C.

**Figure 6- Figure Supplement 1.**
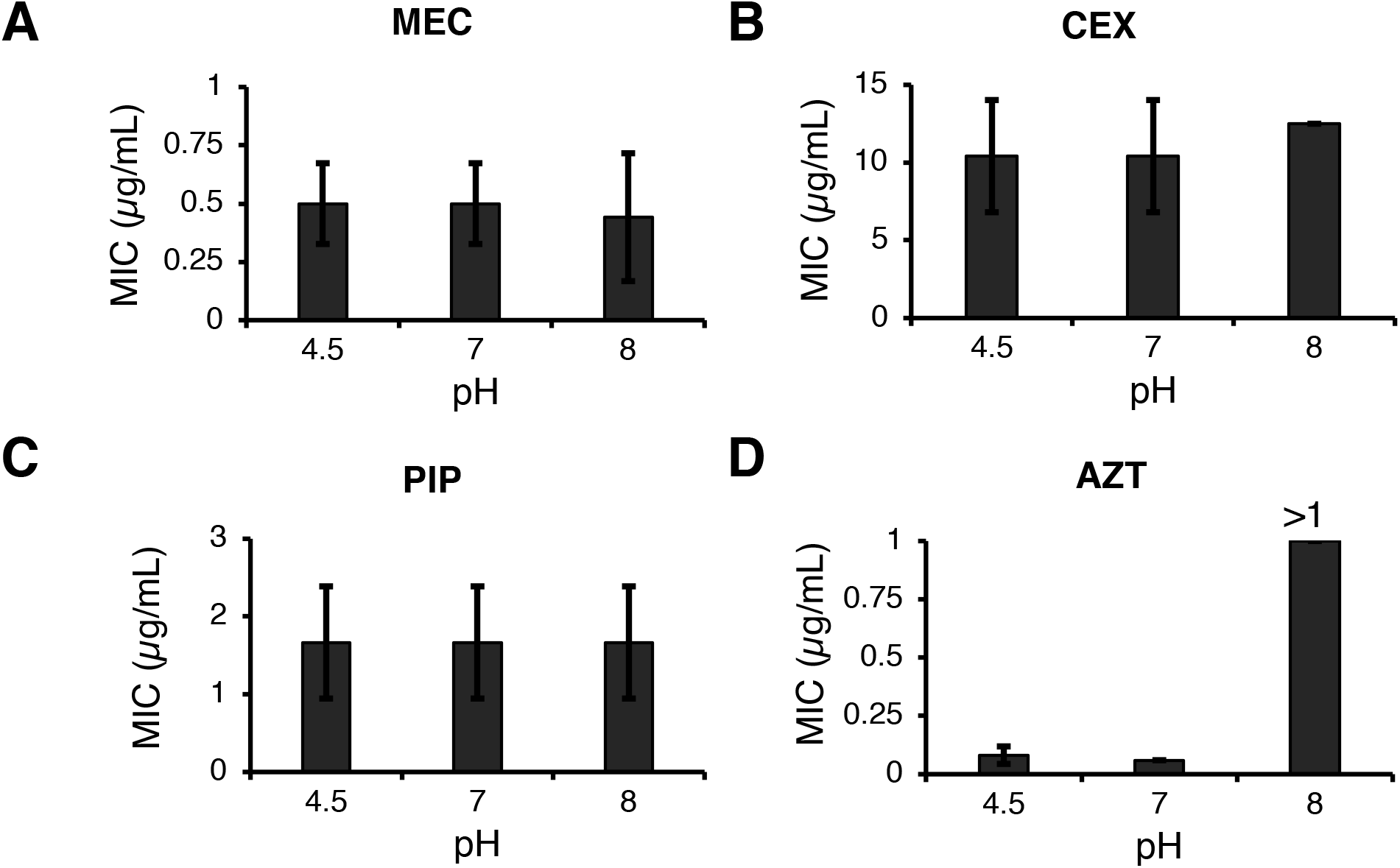
Stability of β-lactam antibiotics across pH values. Mecillinam (A), cephalexin (B), piperacillin (C), and aztreonam (D) were incubated in LB media at pH 4.5, 7.0, or 8.0 for 20 hours then inoculated into microtiter dishes with MG1655 for determination of the minimum inhibitory concentration as previously described. Bars represent average minimum inhibitory concentration +/- standard deviation from at least three independent biological replicates.

**Figure 6- Figure Supplement 2.**
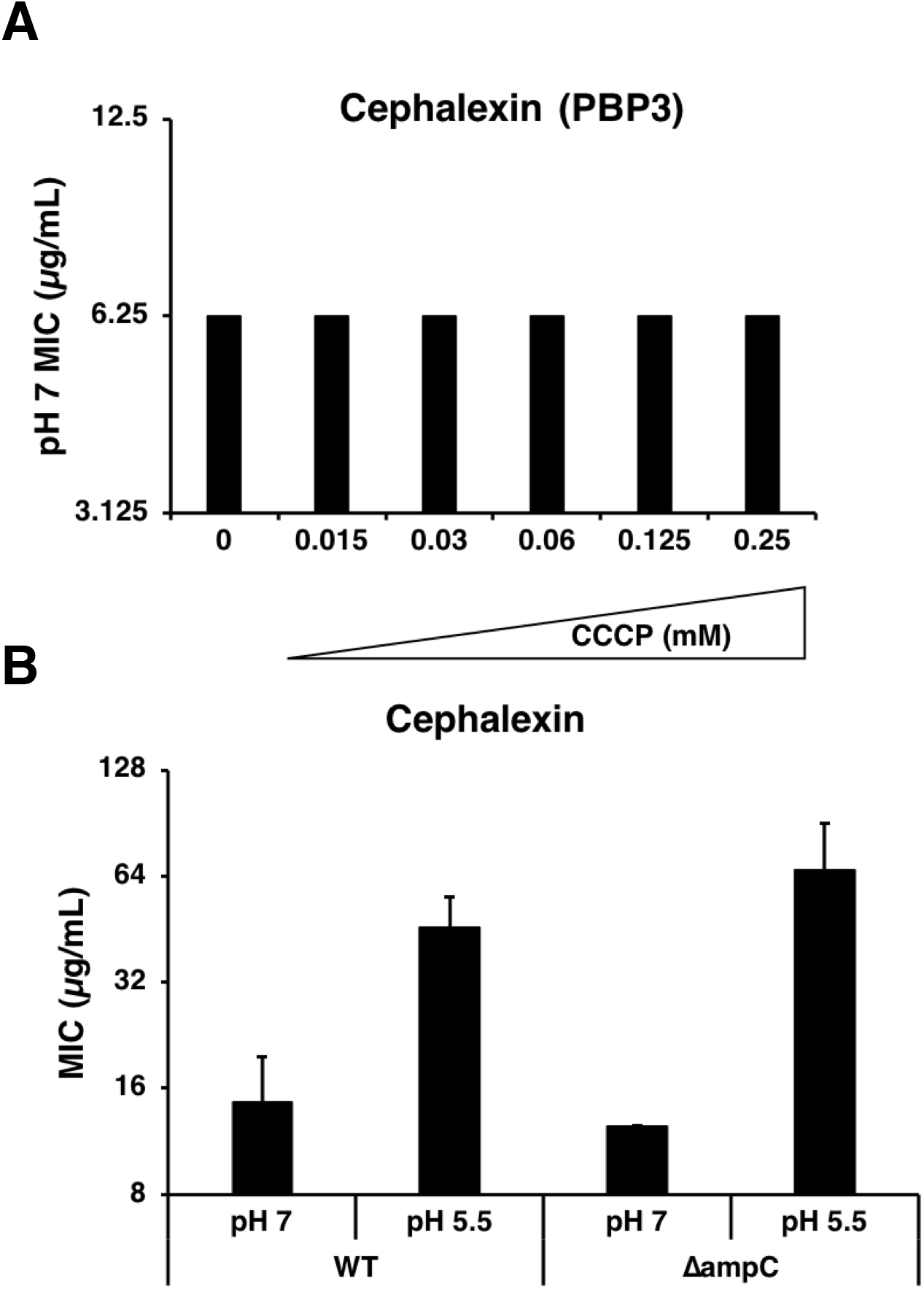
Proton motive force and AmpC β-lactamase do not confer pH-dependent resistance to cephalexin. A) Minimum inhibitory concentration of cephalexin at pH 7.0 in the presence of various concentrations of proton motive force inhibitor carbonyl cyanide-m-chlorophenylhydrazone. B) Comparison of the minimum inhibitory concentration of cephalexin at pH 7.0 and pH 5.5 between wild type and Δ*ampC* cells.

**Figure 6- Figure Supplement 3.**
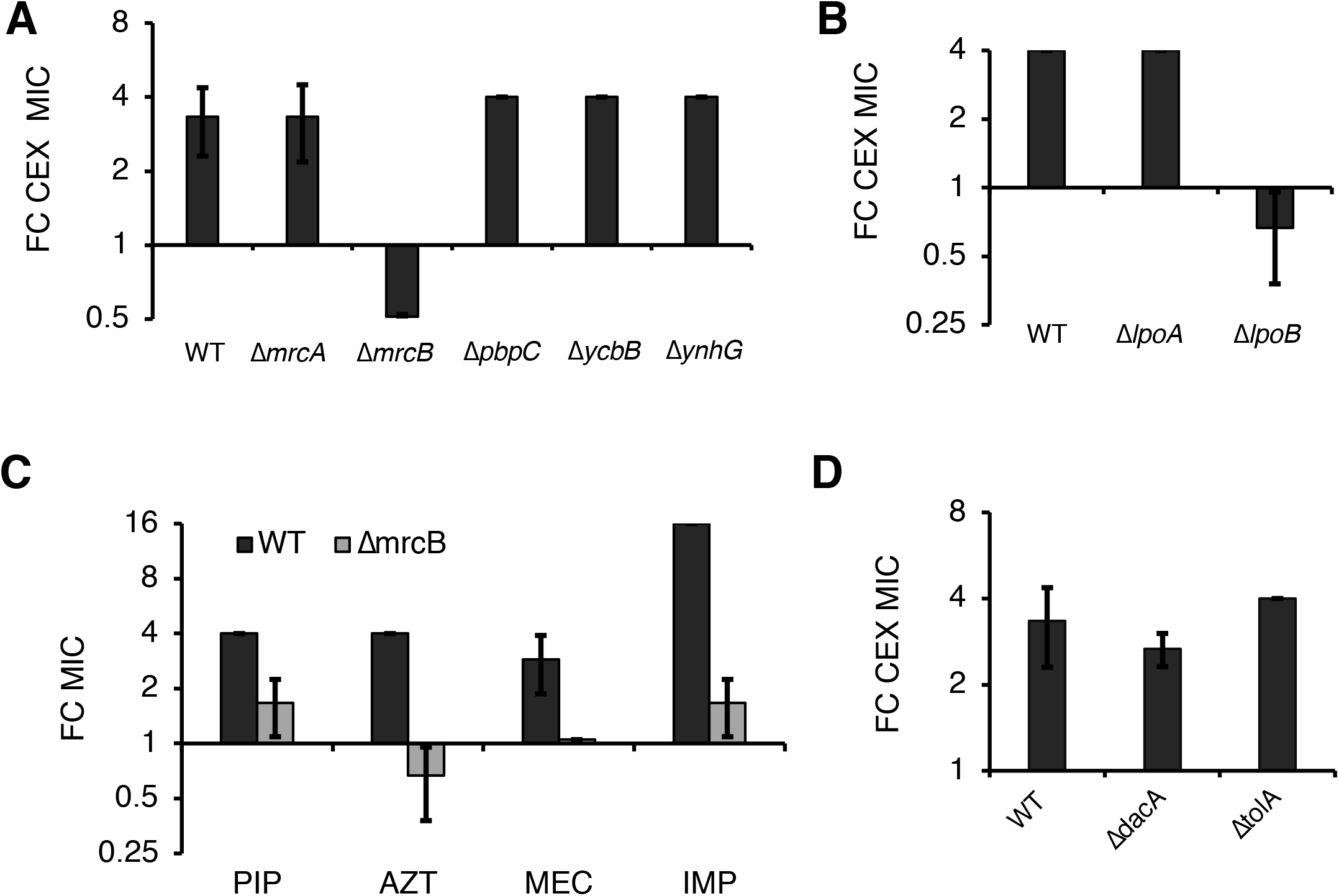
Δ*mrcB* abolishes low pH-dependent resistance independent of growth rate and β-lactam sensitivity. Comparison in the fold change in cephalexin (CEX) minimum inhibitory concentration of mutants in genes encoding nonessential transpeptidases (A) or Lpo lipoprotein activators (B) at pH 5.5 compared to pH 7.0. C) Comparison of the fold change in minimum inhibitory concentration of piperacillin (PIP), aztreonam (AZT), mecillinam (MEC), and imipenem (IMP) for wild type and Δ*mrcB* mutant cells at pH 5.0 compared to pH 7.0. D) Comparison of the fold change in minimum inhibitory concentration of cephalexin (CEX) across wild type, Δ*dacA*, and Δ*tolA* cells. Bars represent average minimum inhibitory concentration +/- standard deviation across three independent biological replicates.

